# Growth control through regulation of insulin-signaling by nutrition-activated steroid hormone

**DOI:** 10.1101/234088

**Authors:** Kurt Buhler, Jason Clements, Mattias Winant, Veerle Vulsteke, Patrick Callaerts

## Abstract

Growth and maturation are coordinated processes in all animals. Integration of internal cues, such as signalling pathways, with external cues such as nutritional status is paramount for an orderly progression of development in function of growth. In *Drosophila*, this coordination involves insulin and steroid signalling, but the mechanisms by which this occurs and how they are coordinated are incompletely understood. We show that production of the bioactive 20-hydroxyecdysone by the enzyme Shade in the fat body is a nutrient-dependent process. We demonstrate that during fed conditions, Shade plays a role in growth regulation, as knockdown of *shade* in the fat body resulted in growth defects and perturbed expression and release of the *Drosophila* insulin-like peptides from the insulin-producing cells (IPCs). We identify the trachea and IPCs as direct targets through which 20-hydroxyecdysone regulates insulin-signaling. The identification of the trachea-dependent regulation of insulin-signaling exposes an important variable that may have been overlooked in other studies focusing on insulin-signaling in *Drosophila*. Finally, we show with IPC-specific manipulations that 20E may both be a growth-promoting and growth-inhibiting signal in the IPCs acting through different nuclear receptors. Our findings provide a potentially conserved, novel mechanism by which nutrition can modulate steroid hormone bioactivation, reveal an important caveat of a commonly used transgenic tool to study IPC function and yield further insights as to how steroid and insulin signalling are coordinated during development to regulate growth and developmental timing.

## Introduction

Growth and maturation are tightly coordinated processes that result in animals of similar, genetically determined and species-specific adult size. Advances in recent years have revealed remarkable similarities in the mechanisms that coordinate growth and developmental timing of maturation in evolutionary distant species such as fly and human with major roles for steroid hormones in maturation and insulin signaling in growth (1-6). Growth and time to maturation universally depend on nutrient availability. With adequate growth and energy storage, maturation is promoted. Likewise, when insufficient energy has been accumulated, antagonistic signals must block onset of developmental transitions to allot more time for feeding, growth, and therefore, energy accumulation (4, 7-12). Throughout development, humoral signals intersect in different tissues to coordinate and balance growth and the onset of developmental transitions (12-15). How this integration occurs and the factors involved, however, are incompletely understood.

The fruit fly *Drosophila melanogaster* is a genetically tractable model to address these questions. In *Drosophila*, juvenile development is marked by a period of exponential growth during the three larval stages or instars (L1-L3), prior to maturation onset. Maturation corresponds to the pupal stage, when the larva metamorphoses into the adult fly and attains its final body and appendage size.

Maturation is regulated in *Drosophila* by the steroid 20-hydroxyecdysone (20E). Its precursor ecdysone (E) is produced in the prothoracic gland (PG) starting from cholesterol. Ecdysone is released into the hemolymph and hydroxylated by the CYP450 enzyme Shade in peripheral tissues to yield the bioactive 20E. The majority of this bioactivation occurs in the *Drosophila* fat body (FB), the functional equivalent of the mammalian liver and adipose tissue (16). 20E is next released and taken up by other tissues and cell types, where it binds a heterodimer of the Ecdysone Receptor (EcR), and Ultraspiracle (Usp) to mediate 20E signalling and transcriptional events via a temporally-defined cascade of downstream nuclear receptors (17,18). Thus, the control of 20E signalling dictates the timing of developmental transitions (19). However, the observations that 20E can inhibit systemic growth but also promote tissue-autonomous growth in the imaginal discs, indicates that the regulation and physiological function of 20E is more complex and diverse depending on tissue, time point and hormone concentration (14, 19-22).

Growth in *Drosophila* is facilitated by the insulin-like peptides (*Drosophila* ILPs, or Dilps), primarily Dilp2, −3 and −5, secreted by the insulin-producing cells (IPCs) in the larval brain (2,3). The Dilps are released into the hemolymph and bind a single insulin receptor (dInR) to activate the highly conserved IIS cascade (1). Reducing systemic IIS either by ablating the IPCs or removing key effectors results in a decreased growth rate, developmental delay and metabolic dysfunction, characterized by a ‘diabetic-like’ phenotype (2,3,23,24). Changes in the growth rate or in growth period length (time to maturation) during larval life result in adults with altered body size (25).

Growth rates and growth period lengths are both highly sensitive to changes in nutritional information. A number of nutrient-sensing mechanisms exist that regulate IIS from the IPCs. Chief among these are nutrient-responsive fat body-derived signals, that are released into the hemolymph in response to nutrients to regulate IIS. Such signals include the recently identified Eiger, Upd2, and Growth-Blocking Peptides which promote or inhibit IIS from the IPCs, respectively (26-28). Both expression and release of all three IPC Dilps are nutritionally-regulated, not only via fat body signals but also through other signals emanating from glia (29) or corpora cardiaca (30). These signals form a complex and dynamic regulatory network which converges on the IPCs and coordinates growth with nutritional status.

This regulation must also be integrated at the level of developmental timing via 20E. Nutrients regulate ecdysone biosynthesis in the prothoracic gland directly - via TOR signalling and control of endoreplication (8,10) - and indirectly - via IPC-derived IIS, which controls both PG size and transcription of the E biosynthesis genes *neverland, spookier, shroud, phantom, disembodied* and *shadow*, collectively known as members of the Halloween Gene family (4,8-10). Reciprocally, peripheral regulation of IIS by 20E has been demonstrated via 20E-sensitive signals such as Dilp6, which inhibits IPC Dilp2 and Dilp5, and 20E-mediated inhibition of *dMyc* (20,21). However, it can be expected that additional layers of regulation as well as additional factors contribute to the coordination of growth and maturation.

Given the fact that (i) the fat body is the central nutrient sensing organ, (ii) *shade* is expressed in the fat body, and (iii) other ecdysone biosynthetic enzymes expressed in the PG are nutrient-sensitive, we hypothesized that *shade* expression in *Drosophila*, and therefore 20E bioactivation, is regulated in a nutrient-dependent manner. We further hypothesized that 20E signalling would be directly required in the IPCs, either to promote IIS as in imaginal discs, or inhibit growth, as via the FB (21).

Here, we show that *shade* expression and 20E synthesis and, thus, bioactivation is nutrient-dependent. During starvation, FB *shade* expression is strongly reduced, and animals are unable to undergo pupation. This failure to undergo pupation was rescued by supplementing steroid hormone, with 20E being more efficient than E. The knockdown of *shade* in the fat body resulted in reduced systemic growth and perturbation of both *dilp* expression and Dilp release from the IPCs. We next showed that ecdysone receptor (EcR) is expressed in the larval IPCs, and when knocked down or otherwise perturbed using the commonly used IPC GAL4 driver line *Dilp2-GAL4^R^* produce extreme growth and metabolic defects reminiscent of starvation. We show that these phenotypes are due to Dilp2 retention and loss of *dilp3* and *dilp5* expression. A detailed analysis of the spatiotemporal expression pattern of *Dilp2-GAL4^R^* revealed that it is not only expressed in the IPCs but also in the trachea. The strong IIS reduction is the combined effect of 20E perturbation in trachea and IPCs, with the most prominent contribution by the trachea. We also provide evidence that the trachea may itself be a source of *dilp2*. Finally, we demonstrate a role for 20E in regulating growth and IIS using IPC specific manipulations. We put forth evidence and propose a model wherein 20E acts as both an IIS-promoting and -inhibiting signal. Nuclear receptors early in the cascade promote IIS, whereas nuclear receptors late in the 20E signalling cascade inhibit IIS, recapitulating the epistatic model demonstrated in 20E regulation of E biosynthesis in the PG and originally proposed by Ashburner in 1974 (31,32).

Our data contribute to our understanding of how growth and maturation are coordinated and we identify three additional regulatory levels at which nutritional cues are integrated in the insulin-steroid regulatory network. It involves key peripheral tissues, such as the fat body and trachea, which produce signals that are integrated at central promoters of growth and maturation, insulin and steroid-producing tissues, respectively, which in turn communicate with one another directly.

## Results

### Conversion of ecdysone to 20-hydroxyecdysone is nutrient-dependent

Ecdysone (E) is synthesized in the PG starting from cholesterol by the consecutive action of enzymes encoded by the Halloween Gene Family (*neverland, spookier, shroud, phantom, disembodied* and *shadow)* (4,8,10). A final Halloween Gene, *shade*, encodes the CYP450 E20-monooxygenase that is expressed in the fat body, the primary nutrient sensing organ, where it converts ecdysone in the bioactive 20-hydroxyecdysone (20E) (Figure 1, A; (16)). We first asked whether *shade* expression is nutrient-sensitive. To test this, we conducted starvation experiments at 72 hours after egg laying (h AEL) and compared *shade* expression between fed and starved larvae 24h later (Figure 1, B). This resulted in a significant reduction of detectable *shade* expression levels in fat body but not gut, another tissue in which *shade* is expressed. A comparable decrease of *shade* transcript levels was observed upon knockdown of the amino acid transporter *slimfast (slif)* in the fat body (Figure 1, C). This indicates that *shade* regulation in the fat body is nutrient-dependent, and amino acid-sensitive.

**Figure 1:**
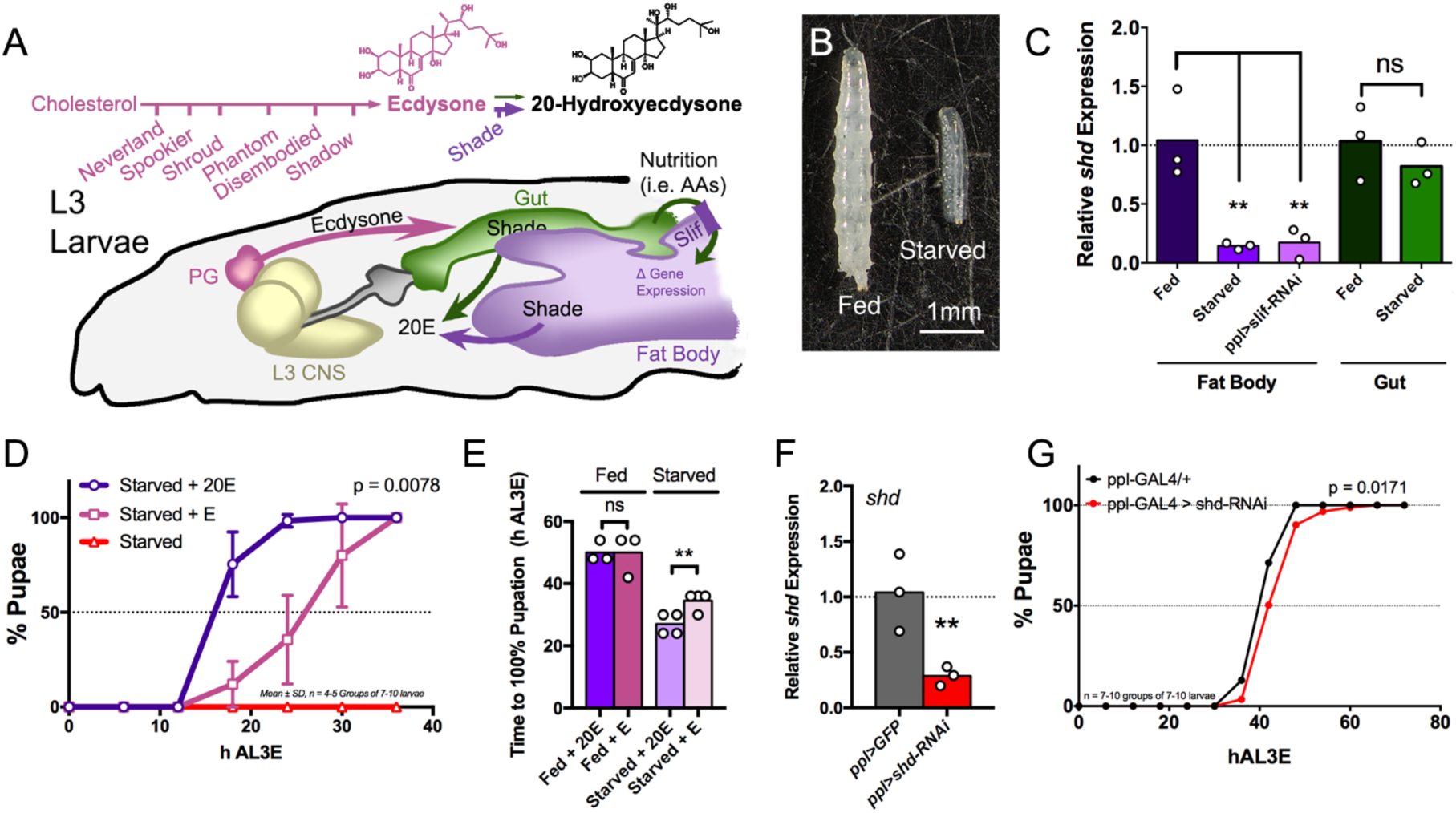
20E bioactivation and *shade* fat body expression are nutrient-dependent. (A) During larval development, cholesterol is converted into Ecdysone (E) in the prothoracic gland (PG) by a cascade of nutrient-sensitive CYP450 enzymes before being released into circulation. Thereafter, E is bioactivated by Shade in the Gut and Fat Body before being re-released as 20E. Nutrients from larval feeding are transported from the gut to the fat body via transporters, like the amino acid transporter Slimfast (Slif). (B) Larvae starved from 72-96h after egg laying (AEL) are shown at 96h AEL. Larvae were staged and collected from yeasted 1 % agar plates, then collected at 72h AEL and transferred to either new, yeasted sucrose plates (Fed) or 1% non-nutritive agar plates (Starved) for 24h. (C) Starving larvae or knocking down the amino acid transporter *slif* resulted in reduced fat body *shade* expression (p < 0.05, ANOVA). Gut *shade* was unaffected by starvation (p > 0.05, unpaired T-test). RNA was collected from fat bodies or gut dissected from larvae after 24h starvation and time-matched fed controls. (D) Starved larvae do not undergo maturation onset, a phenotype rescued by supplementing 20E or E at 1mg/mL (p < 0.05, ANOVA). (E) The difference in maturation induction between 20E and E was observed during starvation, but not during fed conditions. Larvae were collected at 72h AEL and starved on filter paper supplied with either 1mg/mL 20E or 1mg/mL E in 6% EtOH and 6% EtOH alone as negative control. During feeding, 20E or E was supplemented into 50 μL of a 40% yeast paste solution. The time to pupation of surviving larvae is plotted as a percentage of total animals. The time to reach 100% pupation was compared between conditions. (F) Knockdown of *shade* in the fat body resulted in significantly reduced total body *shade* levels in whole pupae. RNA was collected from time-matched pupae and *shade* levels measured as described above using qPCR. (G) Knockdown of *shade* in the fat body resulted in significantly delayed maturation onset by about 6h (p < 0.05, unpaired T-Test). Time to pupation was determined as above.

Next, we hypothesized that in starved animals, E bioactivation to 20E would be reduced. To corroborate this, we tested the potential of 20E and E to rescue maturation onset in starved animals. Normally, animals starved from 72h AEL onward fail to undergo maturation onset, a phenotype that can be rescued by supplementation of 20E hormone (10,18,33). Under fed condition, supplementation of 20E and E induces precocious pupariation (14,34). We observed that E could rescue maturation onset, but much less efficiently than 20E. This contrasts with hormone feeding in fed animals, where no significant differences were observed between 20E and E supplementation in inducing precocious pupariation, consistent with normal conversion of E to 20E under fed conditions (Figure 1, DE). Finally, we knocked down *shade* in the fat body using the *pumpless (ppl)-GAL4* driver, which resulted in significantly reduced total *shade* expression levels and a developmental delay of about 6h in maturation onset (Figure 1, F-G). Taken together, these results demonstrate that *shade* expression, and therefore 20E bioactivation, is nutrient-dependent, thus identifying a novel mechanism to coordinate nutritional status with steroid biosynthesis and developmental timing.

### 20-hydroxyecdysone, *shade* and growth control

We observed that knockdown of *shade* in the fat body resulted in animals with significantly impaired growth rates. This observation was recapitulated with *cg-GAL4*, and was not due to the *shade-RNAi* line alone (Figure 2, A-B). The effect on growth rates of fat body *shade* knockdown was confirmed using a second RNAi line (Figure S1, A, E). This suggested a role for fat body *shade* in regulating not only maturation onset, but also growth.

**Figure 2:**
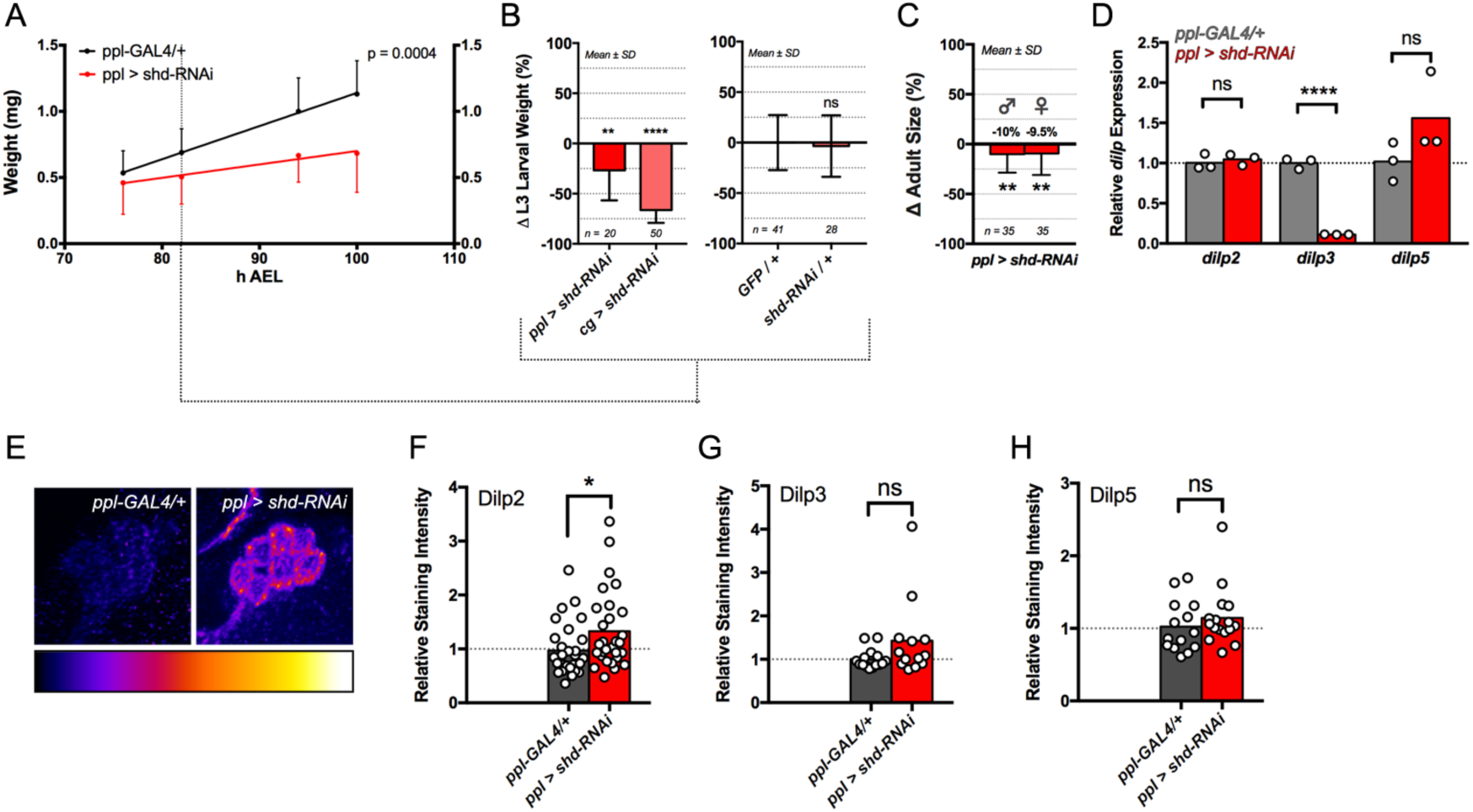
Knockdown of *shade* in the fat body results in reduced larval growth rates and perturbed *dilp* expression and Dilp retention in the brain IPCs. (A) Knockdown of *shade* resulted in reduced larval growth rates compared to the GAL4 alone (mean larval weight ± SD, p = 0.0004, ANCOVA). Collected embryos were distributed at controlled densities of ~50 embryos per agar plate. Larvae were collected at 72h AEL, then weighed every 6 or 12 hours. Raw data points were plotted and a linear regression fit to these raw data. Mean weights are plotted with this linear regression. Slopes of the linear regressions were statistically compared. (B) Knockdown of *shade* using *cg-GAL4* recapitulated the reduced growth phenotype, while the *shade-RNAi* alone did not result in reduced growth. Larvae were weighed at 84h AEL and compared to *cg-GAL4/+* larvae. (C) Eclosed adult *ppl-GAL4 > shade-RNAi* flies were significantly smaller than *ppl-GAL4/+* flies reared in controlled densities as described above. Flies were weighed between 3-7 days old without exposure to CO_2_ to prevent desiccation. (D) Knockdown of *shade* in the fat body reduced *dilp3* but not *dilp2* or *dilp5* expression (p < 0.0001, T-test). RNA was collected from whole *ppl-GAL4 > shade-RNAi* and *ppl-GAL4/+* larvae collected at 84h AEL. (E-H). Knockdown of *shade* in the fat body resulted in retention of Dilp2 (p < 0.05, unpaired T-test). No differences in Dilp3 or Dilp5 staining intensity were observed in *ppl-GAL4>shade-RNAi* compared to *ppl-GAL4/+* IPCs (p > 0.05, unpaired T-test). Larval brains were dissected and fixed, then stained for either Dilp2 & Dilp3 or Dilp2 & Dilp5. Staining intensities were measured in IPC cell bodies using FIJI and plotted relative to *ppl-GAL4/+* controls.

Recent studies have identified a number of non-autonomous fat-body derived signals that regulate IIS and growth via the brain insulin-producing cells (IPCs) (20,26-28). These factors exhibit nutrient-dependent expression in the fat body, and contribute to regulation of either expression or release of the three IPC-derived Dilps, Dilp2, Dilp3 and/or Dilp5. Thus, we tested whether insulin output from the IPCs was compromised in animals with reduced fat body *shade*. These animals exhibited perturbed *dilp* levels, marked by significant reductions in transcript levels of *dilp3* and a retention of Dilp2 in the IPC cell bodies (Figure 2, D-H; Figure S1, A-D; Figure S2, A). Surprisingly, IPC Dilp3 protein levels in the IPCs were not reduced, suggesting that Dilp3 may also be retained.

Upon starvation, *dilp* expression is reduced in the IPCs. We tested whether overexpressing *shade* in the fat body could rescue starvation-induced *dilp* reductions. Surprisingly, during starvation, *shade* overexpression resulted in increased wet - but not dry - weight, suggesting that *shade* overexpression led to larval water retention (Figure S2, B-F). *shade* overexpression in the fat body and the gut of starved larvae did rescue starvation-dependent *dilp3* reductions, but did not substantially affect *dilp2* or *dilp5* levels, suggesting that shade is sufficient for *dilp3* expression during starvation (Figure S2, G-L, P-R). Feeding 20E to starved animals, however, gave a more complex readout with significant increases in *dilp3* and *dilp5*, but a decrease in *dilp2* (Figure S2, M-O). In summary, we show that fat body *shade* regulates larval growth rates and Dilp expression in the IPCs.

### Targeted disruption of 20E signalling results in severe growth and metabolic defects due to reduced IIS

Based on our results, we hypothesized that 20E has a direct effect on IPCs. In a first step, we focused on the ecdysone receptor (EcR) and used immunohistochemistry with a specific antibody to determine whether the EcR is expressed in the larval IPCs. During the larval stages, the IPCs comprise two clusters of seven cells, one in each brain hemisphere, with proximal neurites and a descending axon bundle, which exits the brain via the esophageal foramen (Figure 3, A). Antibodies directed against EcR revealed expression in the IPCs of L3 larvae, which were marked by expression of a *UAS-sytGFP* driven by *Dilp2-GAL4^R^*, a commonly-used *Dilp2-GAL4* line (Figure 3, B-B''') (2,35).

**Figure 3.**
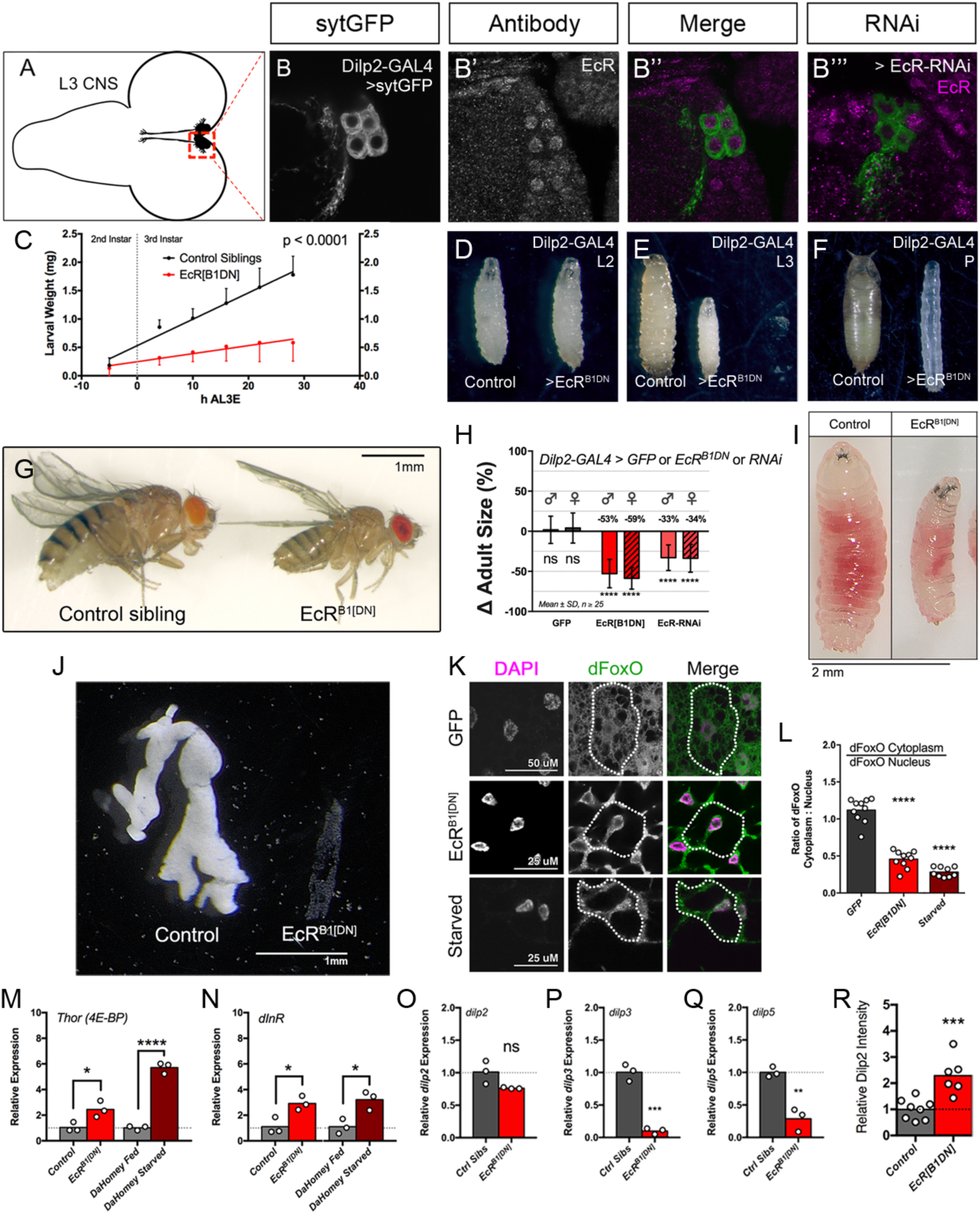
Ecdysone Receptor is expressed in the IPCs, and larvae exhibit severe growth and metabolic defects when EcR signalling is perturbed using *Dilp2-GAL4^R^* due to reduced insulin output from the IPCs, leading to reduced IIS. (A) The insulin-producing cells (IPCs) of *Drosophila* L3 CNS are organized in two clusters of seven neurons (dashed box). (B-B''') Antibody staining for EcR was observed in the IPCs of *Dilp2-GAL4>sytGFP* L3 larvae, and IPC expression was markedly reduced using an *EcR-RNAi*. (C) Larval growth rates were substantially reduced in *Dilp2-GAL4^R^>EcR^B1[DN]^* larvae from late L2 and throughout L3 compared to *EcR^B1[DN]^/CyO* control siblings (mean larval weight ± SD). For growth rate measurement, synchronously timed larvae were collected after L2 ecdysis (~48h AEL) and weights were measured every 6-12 hours. The linear regression was determined with raw weights and is plotted with the mean weight measurements. (D-F) The size phenotype of *Dilp2-GAL4>UAS-EcR^B1[DN]^* larvae has its onset by early third instar (L3) (E). By the end of L3, smaller larvae were leaner and exhibited a clear delay in maturation onset (F; quantified in Figure S3, A). Larvae were staged by 2-3 hour collections on yeasted 1% agar, 20% sucrose plates and compared to control siblings without *Dilp2-GAL4^R^*. Larvae were collected during mid-L1, mid-L2, after L3 ecdysis and at onset of wandering. (G-H) Overexpression of *EcR^B1[DN]^* and genetic knockdown of *EcR* using *Dilp2-GAL4^R^* resulted in smaller adult flies compared to control siblings lacking *Dilp2-GAL4^R^* reared in identical environmental conditions (mean Δ wet weight ± SD, Unpaired T-test, p < 0.05). Expression of a *sytGFP* alone did not affect adult size (H; mean Δ wet weight ± SD, Unpaired T-test, p > 0.05). Adult flies between 3 and 7 days old were not exposed to CO_2_ to prevent desiccation (wet weight) but were instead anaesthetized with chloroform and individually weighed. Expression of *EcR^B1[DN]^* using *Dilp2-GAL4^R^* did not affect feeding (I), yet the same animals displayed a leaner body type and non-autonomous fat body defects, with smaller, transparent fat body tissues compared to control siblings lacking *Dilp2-GAL4^R^* (J). The fat body of a *Dilp2-GAL4^R^>EcR^B1[DN]^* L3 larva and control sibling (*CyO; EcR^B1[DN]^*) is shown (J). (K) The nucleocytoplasmic distribution of dFoxO was changed in these animals with reduced cytoplasmic staining and increased nuclear dFoxO. (L) To quantify dFoxO levels, dFoxO in the cytoplasm was measured relative to the nucleus, then plotted and the cytoplasmic:nuclear dFoxO ratios compared between genotypes. A significant reduction in cytoplasmic dFoxO is observed upon expression of *EcR^B1[DN]^* using *Dilp2-GAL4^R^*, and resulted in significantly increased expression of the dFoxO target genes *thor* (M) and *dInR* (N), comparable to similar phenotypes observed in starved animals. *thor* and *dInR* levels of *Dilp2-GAL4>EcR^B1[DN]^* larvae were compared to respective control siblings. As a positive control, expression of *thor* and *dInR* in *Dahomey w^-^* larvae starved for 24h on a 1 % nonnutritive agar plate were compared to fed larvae of the same genotype. (O-Q) *Dilp2-GAL4^R^>EcR^B1[DN]^* larvae exhibited significant reductions of *dilp3* and *dilp5* (Unpaired T-test, p < 0.05), but not *dilp2* (Unpaired T-test, p > 0.05). (R) Quantification of Dilp2 levels in *Dilp2-GAL4>EcR^B1[DN]^* IPC cell bodies demonstrated a significant retention of Dilp2 compared to respective control siblings. RNA was collected from staged, whole *Dilp2-GAL4>EcR^B1[DN]^* larvae and control siblings reared in identical environmental conditions to control for nutrition and crowding. Transcript levels were normalized to the geometric mean of *Rp49* and *RpS13* expression. For immunohistochemistry, brains were obtained from staged larvae dissected after L3 ecdysis, then stained for Dilp2 together with control siblings. Brains were differentiated by *Dilp2-GAL4^R^*-driven GFP expression and imaged with identical confocal settings. For retention assays, Dilp2 staining of IPC bodies in images was quantified using FIJI. For all experiments, larvae and flies were compared to control siblings to control for genetic background and environmental conditions such as nutrition and larval density.

We next investigated the function of 20E in the IPCs, first disrupting EcR function by using *Dilp2-GAL4^R^* to express a dominant-negative form of the B1 isoform of EcR (EcR^B1[DN]^) in IPCs or to downregulate *EcR* transcript using transgenic RNA interference (RNAi), both of which were previously shown to effectively abolish 20E signalling in peripheral tissues (36,37). Perturbing 20E signalling in the IPCs resulted in significantly reduced larval growth rates from late L2 throughout L3 and a strong delay in maturation onset (Figure 3, C-F; Figure S3, A-B). Animals surviving metamorphosis eclosed into adult flies with a greater than 50% reduction in size compared to control sibling flies reared in identical nutritive and crowding conditions (Figure 3, G-H).

Despite being able to feed and process food normally (Figure 3, I), these larvae also exhibited clear energy storage defects, as exemplified by their near-transparent fat bodies compared to fat bodies taken from control siblings reared in identical environmental conditions (Figure 3, J). Triglyceride levels and lipid content of these fat bodies were substantially reduced (Figure S3, C-D, F). The cells of the fat bodies were also markedly smaller, which, together with the metabolic and growth defects is consistent with a reduced IIS phenotype (Figure S3, E).

Energy homeostasis and lipid metabolism are altered upon reduction in systemic IIS (23,24,38). In *Drosophila*, like in other animals, IIS is thought to promote energy storage and lipid synthesis opposite lipolysis-promoting glucagon-like signals, such as *Drosophila* Adipokinetic Hormone (AKH) (38–41). Hallmarks of reduced IIS include not only reduced energy stores and smaller cells and body size, but also nuclear localization of dFoxO and increased expression of dFoxO targets genes *thor (4E-BP)* and *dInR* (42,43). To further confirm that IIS levels are reduced in *Dilp2-GAL4^R^* > *EcR^B1[DN]^* larvae, we analyzed nuclear localization of dFoxO in their fat bodies and quantified expression of the dFoxO targets *thor* and *dInR*. Staining for dFoxO in fat bodies of these larvae revealed a significant reduction of cytoplasmic dFoxO in *Dilp2-GAL4^R^>EcR^B1[DN]^* fat bodies compared to fat bodies of *Dilp2-GAL4^r^*/+; *sytGFP*/+ controls reared in identical nutritive and crowding conditions, similar to what is observed in starved animals (Figure 3, K-N).

Next, we sought to determine whether IPC output was disrupted in *Dilp2-GAL4^R^* > *EcR^B1[DN]^* animals. We quantified *dilp2, dilp3* and *dilp5* larval transcript levels and staining intensity of Dilp protein in the IPCs. This revealed a clear reduction of *dilp3* and *dilp5* transcript and protein, and while *dilp2* transcript levels were not significantly different than that of control siblings, Dilp2 protein was retained in the cell bodies (Figure 3, O-R; Figure S4). Taken together, these results demonstrate that *Dilp2-GAL4^R^* > *EcR^B1[DN]^* animals exhibit significantly reduced growth and metabolic storage defects due to reduced insulin output from the IPCs, leading to reduced systemic IIS.

### Growth and metabolic phenotypes depend on trachea and IPCs

The results obtained with *Dilp2-GAL4^R^-mediated* perturbation of 20E signalling suggested an exciting, potentially significant contribution of nutrient-dependent 20E signalling to IPC development and function in growth regulation. However, several anomalies led us to revisit these results. Mainly, the observed size phenotypes were comparable to IPC-ablation studies, and more severe than those reported in *Δdilp* mutant studies. Of note, even among these studies, there is a vast difference in the reported phenotypes between groups and publications (2,3,23,24). To further characterize the role of ecdysone signaling in IPCs, we first characterized in detail the spatiotemporal expression pattern of the *Dilp2-GAL4^R^*-driver. Starting from late embryonic stages and throughout larval development, the *Dilp2-GAL4^R^* driver showed expression in the IPCs and salivary glands (Figure S5). Disruption of the salivary glands does not affect IPC development or function (2). We further characterized *Dilp2-GAL4^R^* spatiotemporal activity using a recently described Gal4 technique for real-time and clonal expression (G-TRACE) (44). This works using *Dilp2-GAL4^R^* to drive expression of an NLS-RFP and a recombinase to flip out a STOP cassette between a ubiquitous *p63* promoter and an NLS-GFP. Thus, NLS-RFP expression marks current GAL4 activity, whereas NLS-GFP expression marks transient, past GAL4 activity prior to when samples were collected for imaging (Figure 4, A; (44)). Using this technique, we found that the *Dilp2-GAL4^R^* driver is active not only in the IPCs and salivary glands, but also in the trachea. In all dissected animals, *Dilp2-GAL4^R^* and its corresponding line from the Bloomington *Drosophila* Stock Center (BDSC) *Dilp2-GAL4^37516^* both drove expression strongly in the nuclei of L3 trachea (Figure 4, B-C’). These *Dilp2-GAL4* lines were generated by cloning three copies of an 859 bp fragment upstream of the *dilp2* gene locus in front of GAL4, a construct that was subsequently randomly integrated into the *Drosophila* genome (2)(see Figure 4H for a schematic representation). We observed that tracheal expression is not due to the genomic insertion site of this fragment in these lines, but rather may be intrinsic to the transgene itself, as re-isolating and cloning a single copy of the upstream fragment into a different binary expression system, the *LexA/LexAop* system, also results in tracheal activity of the *Dilp2-LexA* (Figure 4, D-D’). Combining *Dilp2-GAL4^R^* with a *breathless-GAL80 (btl-GAL80)* - a tracheal blocker of GAL4 activity (45) - abolished tracheal activity of the *Dilp2-GAL4^R^*, but retained IPC activity (Figure 4, E-E’). Furthermore, two other *Dilp2-GAL4* lines, *Dilp2-GAL4^215-1-1-1^* (46) and *Dilp2-GAL4^96A08^* (47), cloned from other, single upstream *dilp2* fragments, did not exhibit tracheal activity above low, autofluorescent background (Figure 4, F-G'). Our observation that the *Dilp2-GAL4^R^* line drives expression in IPCs and trachea raised two questions: (i) does the enhancer activity reflect *bona fide* expression of *dilp2* in the trachea, and (ii) what is the relative role of IPCs and trachea in the observed growth phenotype in *Dilp2-GAL4^R^* > *EcR^DN^* animals? To address the first question, we determined *dilp2* transcript levels in trachea. *dilp2* transcript was detected in the trachea of *Dahomey w*-animals, accounting for ~10% of total body *dilp2* expression (Figure 4, I). Taken together, *Dilp2-GAL4^R^* and *Dilp2-GAL4^37516^*, the most commonly used GAL4 lines to transgenically manipulate the IPCs, exhibit strong activity in the trachea that may reflect endogenous *dilp2* expression.

**Figure 4:**
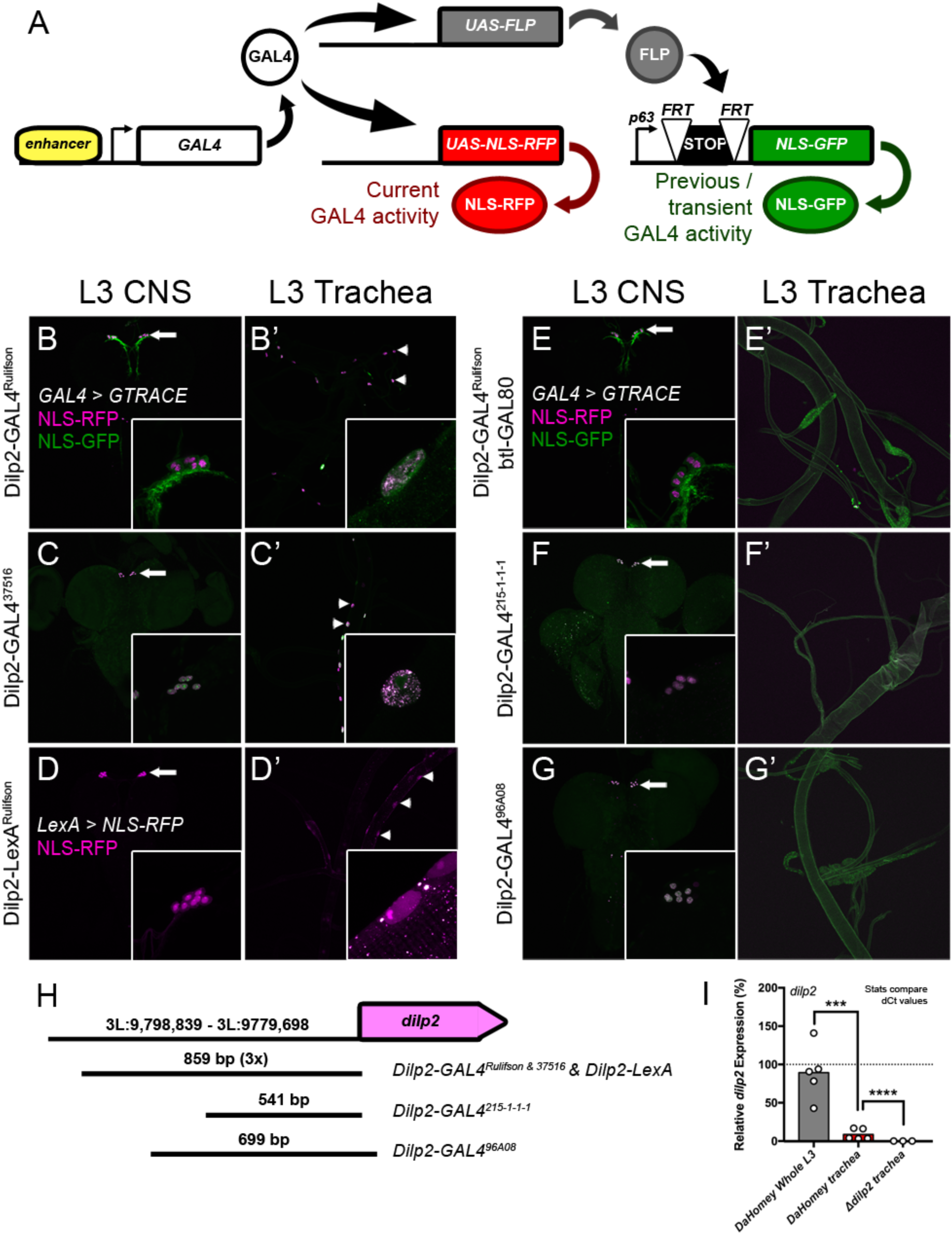
Commonly used *Dilp2-GAL4* drivers exhibit strong tracheal activity that may reflect endogenous *dilp2* expression. (A) The G-TRACE system allows identification of both current and past GAL4 activity. When combined with an enhancer-driven GAL4, the GAL4 drives expression of an NLS-RFP, which marks current activity, and a Flippase (FLP). FLP is a recombinase that will loop and recombine out a STOP cassette in between a ubiquitous p63 promoter and NLS-GFP. Thus, cells with GAL4 activity and all future daughter cells will be GFP-positive, marking transient GAL4 activity. (B-B’) Combining Dilp2-GAL4^R^ with the G-TRACE system results in labelling IPC and tracheal nuclei. IPC soma, neurites and axons are also labelled due to a *UAS-sytGFP* present in the stock. (C-C’) Dilp2-GAL4^37516^, which contains the same insertion as Dilp2-GAL4^R^, also drives expression in both IPCs and trachea. (D-D’) Cloning a single copy of the 859bp enhancer (corresponding to the fragment used as trimer in Dilp2-GAL4^R^) into the LexA/LexAop system resulted in a Dilp2-LexA, which also drove expression of an NLS-RFP in both IPCs and trachea. (E-E’) Tracheal activity of Dilp2-GAL4^R^ is abolished when combined with a btl-GAL80. (F-G’) Both Dilp2-GAL4^215-1-1-1^ and Dilp2-GAL4^96A08^ drive expression in the IPCs, but not the trachea. Trachea are easily visible in E’, F’ and G’ due to autofluorescence. *Dilp2-GAL4>G-TRACE* or *LexA>NLS-GFP* larvae were collected at wandering L3. All tissues were mounted without immunohistochemical staining and imaged. (H) Schematic representation of the Dilp2 regulatory sequences used in the different *Dilp2-GAL4* driver lines. Insertions from common *Dilp2-GAL4* lines. *Dilp2-GAL4^R^* and *Dilp2-GAL4^37516^* consist of 3 copies of a 859 bp dilp2 upstream fragment regulating the GAL4 transcriptional activator, while *Dilp2-LexA* is controlled by a single copy of the upstream fragment. (I) qPCR analysis detects *dilp2* transcript levels in *Dahomey w*- trachea that account for ~10% of total *dilp2* transcript. For qPCR, 50 trachea were dissected from wandering L3 for each biological replicate and *dilp2* expression measured. Normalized ΔCt values were compared. Trachea were taken as *dilp2*- positive if expression was significantly higher than *Δdilp2* deletion mutants, then compared to total *dilp2* expression from wandering, whole *Dahomey* larvae (ANOVA, p < 0.05). The dashed line represents total *dilp2* expression in whole larvae.

To address the second question, we first tested whether perturbing 20E signalling in the trachea can phenocopy growth defects using a tracheal *btl-GAL4*. At 25°C, this condition was lethal, while at 18°C adult flies emerged that exhibited significant growth defects (Figure S6). Next, we repeated experiments with *Dilp2-GAL4^R^; btl-GAL80* and *Dilp2-GAL4^215-1-1-1^*, both of which do not exhibit tracheal GAL4 activity (Figure 4, E’, F’). Remarkably, repeating growth experiments and qPCR measurement of *dilp2, dilp3* and *dilp5* revealed a substantial reduction in the severity of growth defects, and the disappearance of *dilp3* and *dilp5* expression reduction (Figure 5). Flies still exhibited small growth reductions (Figure 5, B), though overexpressing *EcR^B1[DN]^* in the IPCs with *Dilp2-GAL4^215-1-1-1^* resulted in no observable size phenotypes (Figure 5, G). These differences in size phenotypes may reflect differences in GAL4 strength, given that *Dilp2-GAL4^R^; btl-GAL80* has 3 copies of the enhancer sequences driving GAL4, while only one copy is present in *Dilp2-GAL4^251-1-1-1^*. Nonetheless, measuring circulating Dilp2 in these animals demonstrated a 20% reduction in secreted Dilp2 protein, suggesting that 20E may function in the IPCs to regulate IIS, though not at the level of *dilp* transcription, but only Dilp2 secretion (Figure 5, C-F, H-K).

**Figure 5:**
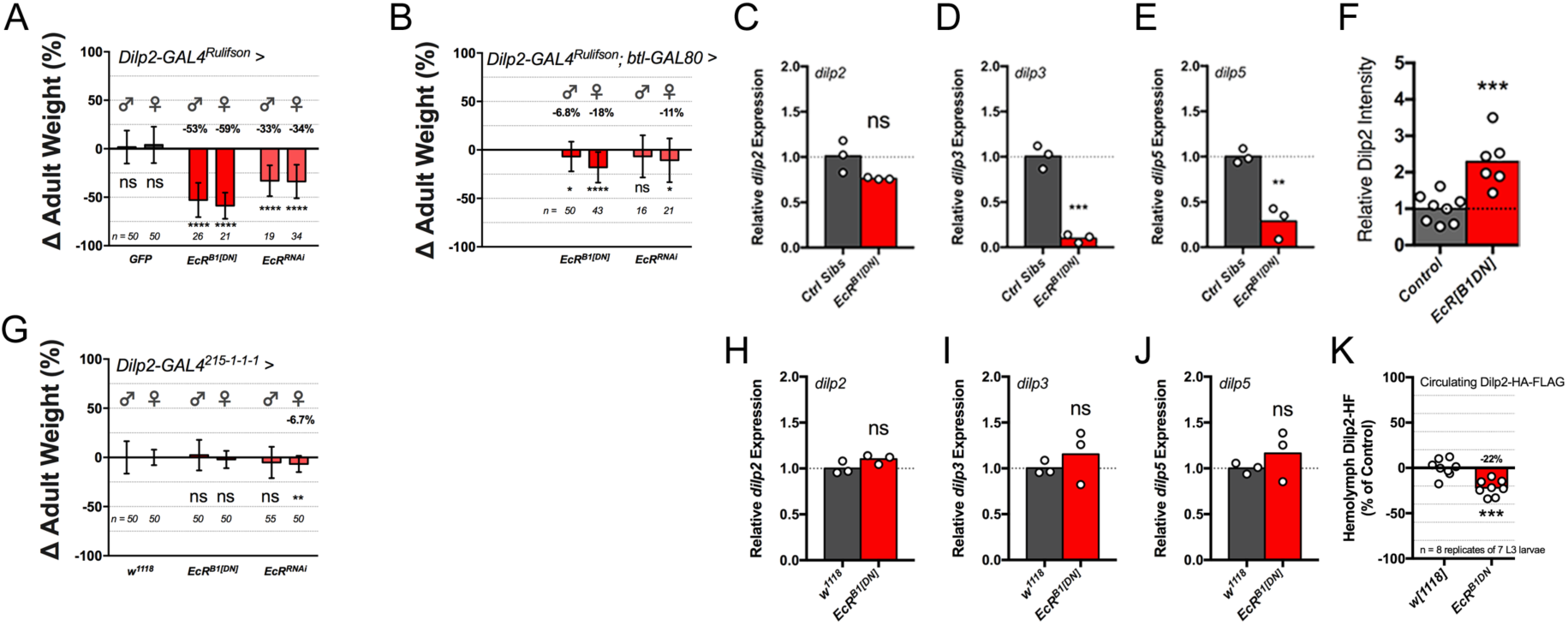
Tracheal perturbations account for most of the growth defects observed in *Dilp2-GAL4^R^*>*EcR^B1[DN]^* animals. (A) Overexpression of EcR^B1[DN]^ or genetic depletion of EcR using *Dilp2-GAL4^R^* alone results in significant growth defects in males and females. Overexpression of EcR^B1[DN]^ or genetic depletion of EcR using (B) *btl-GAL80* or (G) *Dilp2-GAL4^215-1-1-1^* does not or only partially recapitulate the growth defects observed with *Dilp2-GAL4^R^* alone. Knockdown of *EcR* results in significantly smaller females with both drivers, while overexpression of *EcR^B1[DN]^* results in smaller males and females only with *Dilp2-GAL4^R^; btl-GAL80* (unpaired T-test, p < 0.05; mean Δ wet weight ± SD). *Dilp2-GAL4^215-1-1-1^* > *DN* or *RNAi* flies were compared to *Dilp2-GAL4^215-1-1-1^/+* crossed into an identical genetic background (*w^1118^*) and placed at identical densities on fly food to control for crowding. Adult flies between 3 and 7 days old were not exposed to CO_2_ to prevent desiccation (wet weight) but were instead anaesthetized with chloroform and individually weighed. (C-E, H-J) Loss of *dilp3* and *dilp5* transcript levels result mainly from *Dilp2-GAL4^R^-driven* perturbation of *EcR* in the trachea, and not the IPCs. Perturbing 20E signalling specifically in the IPCs using *Dilp2-GAL4^215215-1-1-1^* did not result in any *dilp* transcriptional changes (p > 0.05, ANOVA). RNA was collected from replicates of 7 larvae. For Dilp2-GAL4^215-1-1-1^, *dilp* expression was compared to *Dilp2-GAL4*^215^'^1^'^1^'^1^/+ larvae in the same genetic background as the *UAS-EcR^B1[DN]^* and *UAS-RNAi* lines (*w^1118^)*. Transcript levels were normalized using the geometric mean of *RpS13* and *Rp49* expression. (F) Dilp2-GAL4^R^-mediated overexpression of *EcR^B1[DN]^* resulted in Dilp2 retention in the IPC bodies. Dilp2 staining of IPC bodies in images was quantified using FIJI. For all experiments, larvae and flies were compared to control siblings to control for genetic background and environmental conditions such as nutrition and larval density. (K) Overexpression of *EcR^B1[DN]^* specifically in IPCs using *Dilp2-GAL4^215215-1-1-1^* resulted in a 20% reduction in secreted Dilp2 (p < 0.05, ANOVA). 1 μL of hemolymph was collected from 8 replicates of 7 larvae and circulating Dilp2 levels were measured by using an ELISA assay where Dilp2 with HA and C-terminal FLAG tags was affixed to plates coated with anti-FLAG and detected using anti-HA-peroxidase as described in (46).

In summary, the impact of 20E on growth is mediated via trachea and IPCs. The effect of the trachea is reminiscent of the previously described impact of hypoxia on IPC function leading to impaired growth and lipid metabolism via Dilp2 retention and *dilp3* and *dilp5* expression reduction (48).

### Nuclear receptors differentially regulate IIS in IPCs

To confirm and elaborate on the role of 20E-signalling in the IPCs, we determined whether 20E-signalling components are expressed in the larval IPCs with available antibodies for the typical 20E nuclear receptors, other than EcR (17). The heterodimerization partner of EcR, Ultraspiracle (Usp), as well as other nuclear receptors Ecdysone-induced protein 75B (E75), *Drosophila* Hormone Receptor 3 (DHR3) and Ftz transcription factor 1 (Ftz-f1) were also detected in the IPCs during L3. All of these nuclear receptors could be efficiently genetically depleted in the IPCs with exception of Usp, for which knockdown was incomplete. The latter is consistent with an earlier report that Usp expression is difficult to reduce in the CNS (Figure 6, A-D; (49)).

**Figure 6:**
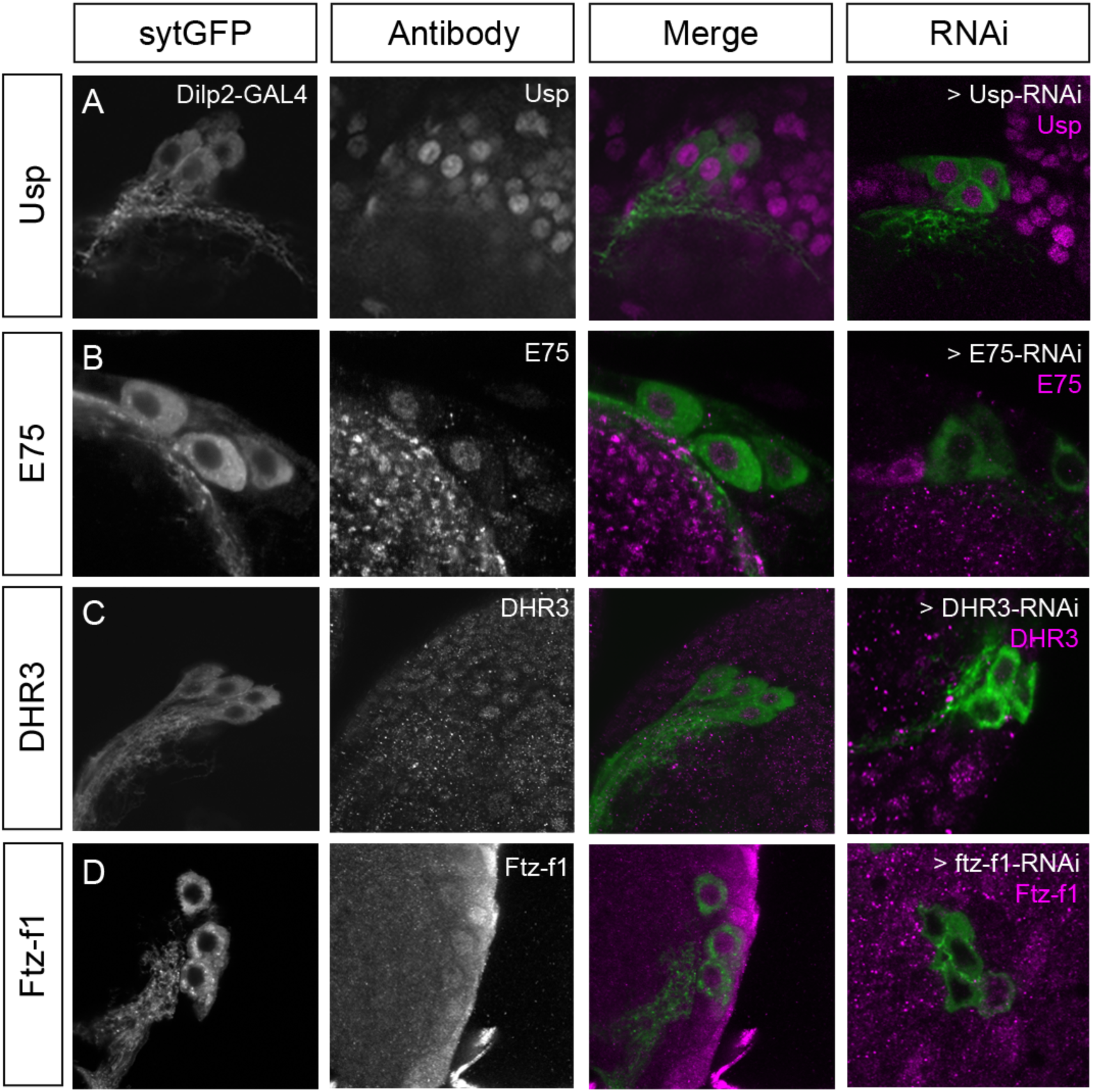
Typical 20E nuclear receptors are expressed in the larval IPCs. (A) Usp expression could be detected and reduced in the IPCs, though knockdown was incomplete. (B-D) E75, DHR3 and Ftz-f1 expression were all detected in the IPCs of L3 brains, and could be effectively depleted using transgenic RNAi-mediated knockdown. For antibody staining, IPCs were marked by *Dilp2-GAL4^R^* driving *syt-GFP*. Brains were collected from L3 larvae, fixed and then stained with individual antibodies directed against the various nuclear receptors. For antibody concentrations and origins, see Material & Methods.

Next, we tested whether genetic depletion of these nuclear receptors in the IPCs resulted in growth or IIS defects using all three GAL4 driver combinations (Figure 7). First, with *Dilp2-GAL4^R^*, knockdown of all nuclear receptors except *DHR4* resulted in severe growth and IPC morphology defects and reductions in *dilp* transcription reminiscent of those observed with perturbations of *EcR* (Figure 7, A, D-F; Figure S7, S8). Knockdown of *ftz-f1* with *Dilp2-GAL4^R^* resulted in growth defects comparable to *EcR^B1[DN]^*, and resulted in similar Dilp2 retention (Figure 7, H). These phenotypes also depend on trachea and IPCs and were drastically reduced - or, as with *ftz-f1* and *DHR3* - disappeared or even reversed (Figure 7, B-C, H-K) when restricted to IPCs. When manipulating 20E nuclear receptors in IPCs only by using both *Dilp2-GAL4^R^; btl-GAL80* and *Dilp2-GAL4*^215-1-1-1^, we observe a significant size reduction with knockdown of *E75* in both males and females, similar to as when knocked down with *Dilp2-GAL4^R^*. Knockdown of *ftz-f1* produced no clear effect on size - a reduction in *Dilp2-GAL4^R^; btl-GAL80* > *ftz-f1-RNAi* males, but an increase in *Dilp2-GAL4^215-1-1-1^*> *ftz-f1-RNAi* males. Surprisingly, knockdown of *DHR3* resulted in larger flies with both drivers when genetically depleted specifically in the IPCs, suggesting that the trachea effect masked the true phenotype entirely. Next, knockdown of *DHR4* specifically in the IPCs resulted in smaller females with both drivers, but no effect on males. Finally, knockdown of Usp only resulted in smaller flies with *Dilp2-GAL4^215-1-1-1^* (Figure 7, B-C). Measuring *dilp* transcript levels of *Dilp2-GAL4^215-1-1-1^* > *NR-RNAi* larvae revealed no reductions in *dilp3* and *dilp5* transcript levels. However, *dilp2* transcription was reduced with knockdown of *DHR3* and *DHR4*, which also saw significant increases in *dilp5*. Knockdown of *DHR4* and *Usp* also resulted in a significant increase in *dilp3* transcription (Figure 7, H-J). Finally, measuring circulating Dilp2 in these animals demonstrated a slight, yet not statistically significant increase in circulating Dilp2 when ftz-f1 was knocked down in the IPCs, again the opposite of what was originally observed in Dilp2 retention with *Dilp2-GAL4^R^*-mediated knockdown (Figure 7, K). Overall, the effects of the nuclear receptors on IIS regulation seem to follow a trend consistent with their known epistatic relationships. Normally, 20E binds a heterodimer of EcR and Usp, which activate early genes such as *E75* and, later, *DHR4*. Late genes such as *DHR3* are activated, and DHR3 protein activates *ftz-f1*, a process that is inhibited by E75. Finally, Ftz-f1 protein inhibits *EcR* in a negative feedback response thought to terminate the potent and diverse effects of 20E signalling (Figure 7, L). Our data suggest that the 20E receptors EcR and Usp promote IIS in the IPCs via E75, which may integrate other signalling pathways like nitric oxide signaling, as well as via the later expressed DHR4. However, the data also suggests that the late 20E nuclear receptors DHR3 and Ftz-f1 serve to inhibit IIS and growth. 20E thus may act both as a signal to promote and to inhibit IIS via temporal activation of these downstream nuclear receptors.

**Figure 7:**
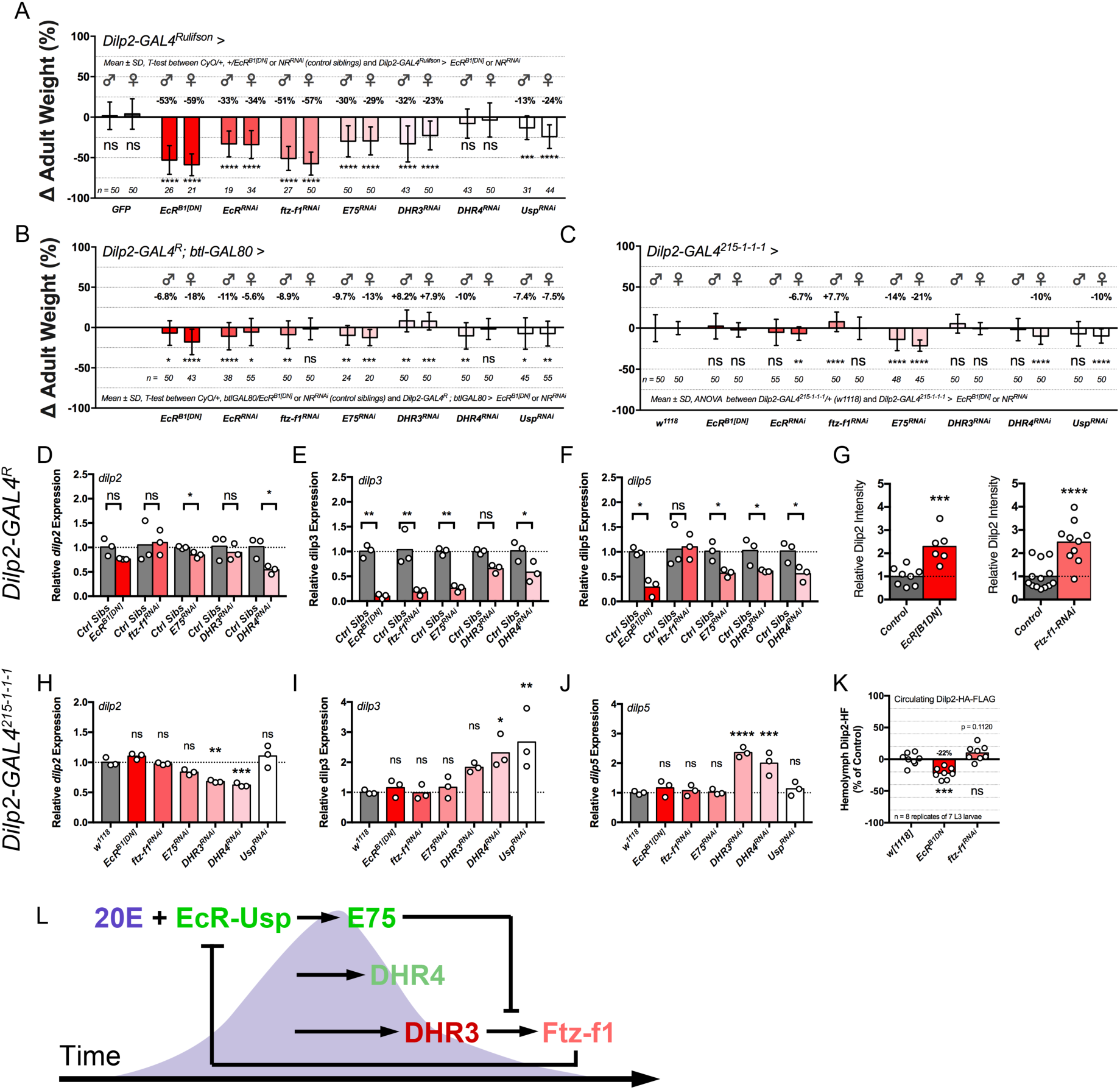
20E is required in both the trachea and the IPCs, wherein individual nuclear receptors have either a positive or negative regulatory role in controlling IIS from the IPCs. (A) Knockdown of 20E nuclear receptors using *Dilp2-GAL4^R^* results in strong size reductions comparable to perturbation of EcR (p < 0.05, unpaired T-test between *EcR^B1[DN]^* or *NR-RNAi/CyO* control siblings and *Dilp2-GAL4^R^*>*EcR^B1[DN1^* or *NR-RNAi*). (B) Blocking tracheal activity of *Dilp2-GAL4^R^* using *btl-GAL80* and conducting IPC-specific nuclear receptor manipulations reduces or even reverses phenotypes. Knockdown of *EcR, E75* and *DHR4* resulted in smaller flies, while knockdown of *DHR3* resulted in larger flies, the opposite of what was observed with the *Dilp2-GAL4^R^* driver alone. (C) IPC-specific perturbations of 20E nuclear receptors using a second *Dilp2-GAL4* driver, *Dilp2-GAL4^215-1-1-1^*, produced comparable results to *Dilp2-GAL4^R^*; btl-GAL80-mediated knockdown. However, knockdown of *DHR4* and *Usp* resulted in only smaller females while knockdown of *DHR3* and *Ftz-f1* produced larger males, but no difference in females was observed. Flies were weighed 3-7 days after eclosion. (D-F; H-J) Knockdown of nuclear receptors using *Dilp2-GAL4^R^* drastically reduced *dilp* expression, but repeating qPCR experiments with *Dilp2-GAL4^215-1-1-1^* did not recapitulate any of these effects. When nuclear receptors were specifically disrupted in the IPCs, there was a reduction in *dilp2* for knockdown of *DHR3* and *DHR4*, an increase in *dilp3* for knockdown of *DHR4* and *Usp*, and an increase in *dilp5* for knockdown of *DHR3* and *DHR4*. (G, K) Perturbing EcR or Ftz-f1 results in Dilp2 retention, and while overexpressing *EcR^B1[DN]^* using *Dilp2-GAL4^215-1-1-1^* significantly reduced circulating Dilp2 levels (p < 0.05, ANOVA), knockdown of *ftz-f1* resulted in a small, but not statistically significant increase (p = 0.1120, ANOVA). Circulating Dilp2-HA-FLAG was detected using ELISA with 1 μL samples of hemolymph collected from replicates of 7 larvae. (L) 20E nuclear receptor epistasis appears to be reflected in the IPCs. Nuclear receptors higher up in the 20E signalling hierarchy appear to have a positive role in IIS regulation, whereas those lower in the 20E signalling have inhibitory roles in IIS regulation.

Taken together, these data show that production of the bioactive 20E is nutrient-dependent, and that 20E nuclear receptors are required in both the trachea and IPCs. Perturbation of 20E signalling in the trachea results in a starvation-like phenotype, inhibiting insulin output from the IPCs and resulting in severe growth and metabolic storage defects. On the other hand, 20E signalling in the IPCs also directly regulates IIS, albeit less prominently than when 20E is also perturbed in the trachea. Our data provide a potentially novel mechanism by which nutrients regulate the production of steroids that act on different tissues, i.e. trachea and IPCs, to control growth and maturation in a complex, intercommunicating network (Figure 8). Finally, we identify the trachea as a tissue with an important regulatory role in growth and maturation. This fact will have to be considered in other studies given the widespread use of the *Dilp2-GAL4^R^* driver line when studying growth and maturation and its regulation.

**Figure 8:**
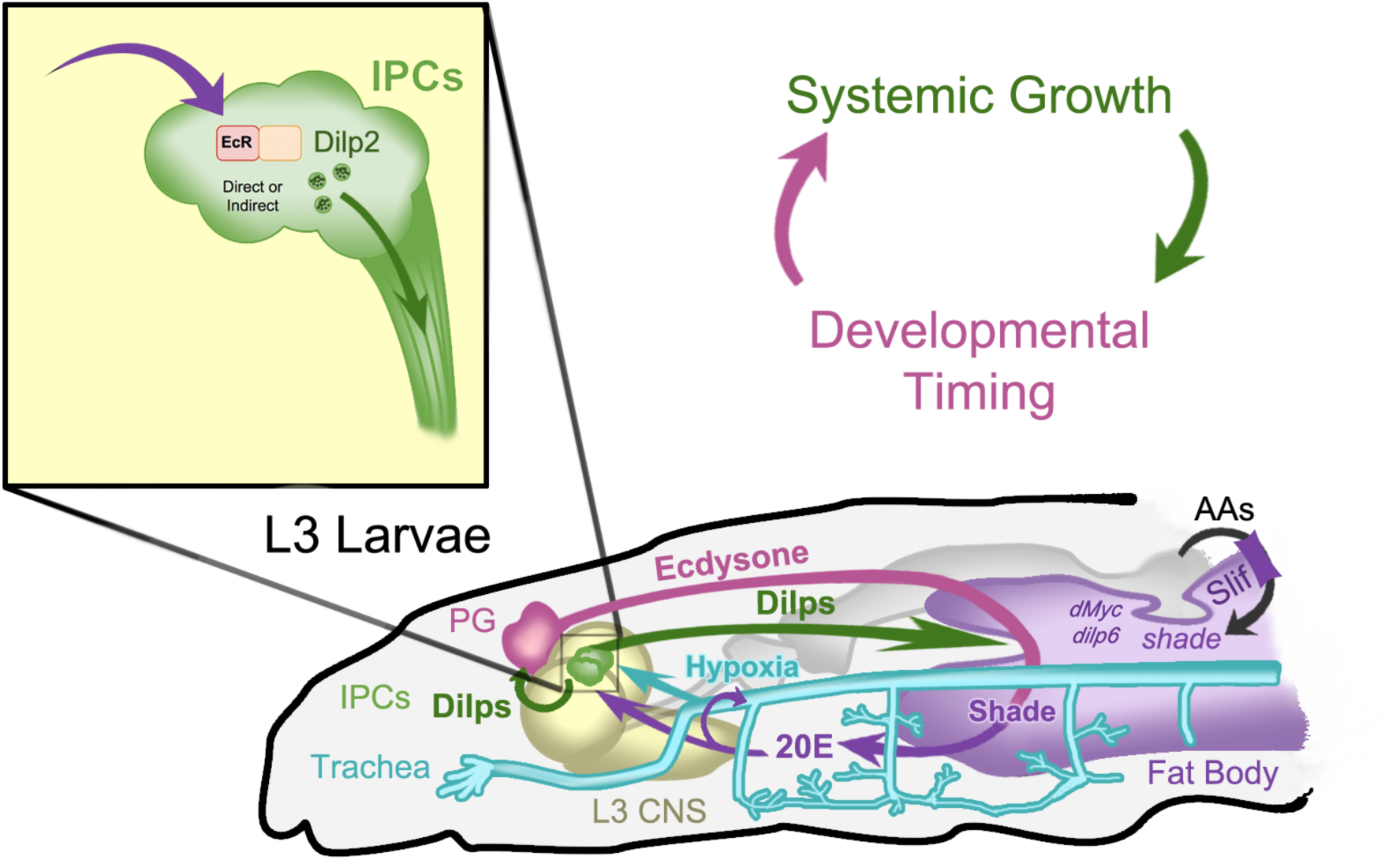
Convergence of humoral signals in central and peripheral tissues maintain an endocrine “goldilocks zone” to coordinate developmental progression and systemic growth. Ecdysone and insulin signalling operate in a double-feedback loop to facilitate normal growth and development that integrates numerous tissues. Ecdysone is produced by the Prothoracic Gland (PG) and converted in the fat body to 20-hydroxyecdysone (20E) by Shade, the expression of which is regulated by nutritional information. 20E contributes IIS regulation via peripheral tissues like the FB and trachea and also via central regulation in the IPCs. IIS can in turn promote ecdysone biosynthesis in the PG (10). Together, 20E and IIS can converge in peripheral tissues or work independently to propagate messages of systemic growth (IIS) and developmental progression (20E) and maintain an endocrine “goldilocks zone” between these two physiological processes.

## Discussion

Integration of growth and developmental timing is an essential requirement for the orderly progression of development. In insects, insulin- and ecdysone signalling regulate the rate and timing of growth in response to nutrition, but the mechanisms by which this occurs and how they are coordinated are incompletely understood. Here, we show that production of 20-hydroxyecdysone by the enzyme Shade in the fat body is nutrient-dependent, that *shade* is required for growth and provide two new avenues by which 20E can regulate IIS-dependent growth - via the trachea and the IPCs. Therefore, our data support the existence of an essential mechanism in the coordination of growth control with developmental progression, and yield important insights as to how nutrition impinges on physiology and development.

To coordinate the onset of developmental transitions with nutritional status, *Drosophila* employs a number of non-autonomous signals that relay nutritional information in feeding larvae from the fat body (FB), the primary nutrient sensor, to the IPCs that regulate IIS (20, 26-28). This nutrient-dependent IIS regulation serves to control both growth and developmental timing, as IPC-released Dilps also promote ecdysone biosynthesis in the PG. Ecdysone biosynthesis can also be nutritionally regulated via TOR signalling in the PG, which, similar to IIS, promotes expression of the enzymes required for synthesis of ecdysone starting from cholesterol. A recent study has shown that this seems to be controlled in part by a TOR-mediated cell cycle checkpoint, with TOR being required for endoreplication of PG cells and production of appropriate levels of ecdysone, that is then converted in the fat body to 20E. In starved animals, levels of E biosynthetic enzymes in the PG and 20E levels are significantly reduced. These animals fail to undergo maturation onset. This can be rescued with supplementation of ingestible 20E, which induces pupation (10). In our work, we show that expression of *shade*, the final enzyme in 20E biosynthesis that is expressed in the fat body, is also reduced upon starvation or knockdown of the amino acid transporter, *slimfast*, and that it can be rescued by supplementation of 20E and, albeit less efficiently, E. We propose that the endoreplication in the prothoracic gland is a mechanism to increase bulk ecdysone production, and that the nutrient-sensitive conversion to 20-hydroxyecdysone in the fat body is a regulatory level that allows rapid adjustment of available bioactive steroid levels to environmental conditions.

The demonstration of nutrition-dependent ecdysone activation draws an intriguing parallel to steroid biosynthesis in mammals. The cholesterol-derived mammalian steroid estradiol is required in both sexes for β-cell development and function in the pancreas. Estradiol can be converted from testosterone by the CYP450 CYP19, or Aromatase, which exhibits expression in a wide range of tissues including the liver and adipose tissues, comparable to the insect fat body (50). Interestingly, evidence in literature points to CYP19 also being nutritionally regulated. CYP19-mediated conversion of testosterone to estradiol is reduced in starved rodents (51), and obese patients have both increased estradiol and measured testosterone to estradiol conversion (52,53). This may imply that a similar nutrition-mediated steroid activation may function in regulating insulin output of mammals, and warrants further investigation particularly in the context of several human disorders with nutrition-associated phenotypes (e.g. obesity and anorexia nervosa) that also impact timing of sexual maturity (5,6).

20E plays a diverse and complex role in growth regulation that differs depending on tissue, time and hormone titer. For example, in the fat body, 20E can both inhibit and promote systemic growth via negative regulation of *dMyc* and positive regulation of *dilp6* (21). Dilp6 released from the FB inhibits Dilp2 and Dilp5 secretion during feeding, but promotes growth during nonfeeding stages (20,54). Another example is in the imaginal discs, where 20E promotes tissue-autonomous growth, though at higher titers during maturation facilitates differentiation of the discs into their prospective adult tissues (14,19). The apparent role of 20E in growth regulation distinct from its traditional role in promoting developmental transitions provides a further layer of complexity. In previous work, it was shown that transgenic manipulations during L3 of 20E titers via IIS inhibition in the PG altered systemic growth without affecting developmental timing (4). In our work, we show that knockdown of the 20E biosynthesis enzyme *shade* in the FB reduced both growth and delayed maturation onset. Interestingly, FB *shade* has been previously implicated in developmental timing, though a role in growth has not been described and, together with our other results, suggests this regulation is a nutrient-dependent process. We also performed reciprocal experiments to see if overexpressing *shade* in the fat body during nutrient deprivation affected body size and *dilp* levels. During starvation, overexpressing *shade* rescued nutrient-dependent reductions of *dilp3*. We also observed a weight gain in these animals, an observation we were able to attribute to water retention. This draws a further interesting parallel to mammals, where exogenous steroid application similarly results in increased water retention (55). Given the previously identified connections between the FB and IPC-regulated growth, we wondered whether insulin output from the IPCs was affected in shade-knockdown animals. Knockdown of *shade* resulted in reduced *dilp3* expression and retention of Dilp2 and potentially also Dilp3. This led us to investigate a possible direct regulatory role of 20E in the IPCs.

Our initial experimentation identified phenotypes of extreme growth and IIS reduction when 20E was perturbed in the IPCs using *Dilp2-GAL4^R^*, the most commonly used *Dilp2-GAL4* driver. We then mapped in detail the spatiotemporal expression pattern generated by this GAL4 driver and found expression in the trachea in addition to the well-known expression in the IPCs. A significant part of the observed phenotypes was next shown to depend on ecdysone signaling in the trachea, thereby identifying the trachea as an important organ in the regulation of growth and maturation. Given that the *Dilp2-GAL4^R^* driver has been used in numerous studies (>70) on the regulation of growth and maturation, it will be important to carefully revisit whether the published observations are not due to tracheal activity of the driver and, consequently, genetic manipulations perturbing tracheal development and function. Revisiting some of these studies with the *Dilp2-GAL4^R^; btl-GAL80* or a second *Dilp2-GAL4* driver may also contribute to resolving some of the contradictions that exist between published observations. A notable example of this is differences in phenotypic severity between the various IPC ablation studies, which show a range of size reductions from more than 50% to half that dependent on the *Dilp2-GAL4* lines that were used (2,3). In summary, these observations emphasize the necessity for performing tissue-specific manipulation experiments with multiple drivers, and raise concerns over reproducibility with respect to conclusions drawn from experiments made with single GAL4 drivers.

We identify an important role of 20-hydroxyecdysone in the third instar larval trachea to regulate growth and maturation. Previously, 20E had only been implicated in tracheal morphogenesis during embryonic development (56). When perturbing 20E signalling using *Dilp2-GAL4^R^*, IIS phenotypes only arose during L3 (Figure 3, D-G). These phenotypes were identical to those previously reported for hypoxic animals, i.e. reduced *dilp3* and *dilp5* transcription and a retention of Dilp2 (48). The fact that the strong growth and metabolic phenotypes disappeared upon blocking tracheal *Dilp2-GAL4* activity, shows that 20E is required in the L3 trachea. One possible explanation is that 20E impacts tracheal function or morphology thereby resulting in hypoxia. This would be consistent with hypoxia being a key regulator of size determination and regulating developmental timing during L3 (48).

Our observation that *dilp2* transcript in the trachea accounted for approximately 10% of total body *dilp2* expression in wandering L3 larvae suggests an alternative explanation, namely that 20E directly regulates dilp2 in the trachea and thereby insulin signaling and growth. However, it remains to be determined whether Dilp2 is secreted from the trachea. Nevertheless, our finding highlights an understudied aspect of growth regulation, namely whether Dilp2, Dilp3 and Dilp5, the Ilps produced by the IPCs, are also produced in and released by other tissues. Functional redundancy and compensatory regulation among the *dilps* have obfuscated identification of these thus far, though future studies using more defined and efficient tissue- and gene-specific manipulations could better elaborate on their contribution to growth.

By blocking tracheal expression of *Dilp2-GAL4^R^* and repeating experiments with a second *Dilp2-GAL4* driver, *Dilp2-GAL4^215-1-1-1^*, we provide evidence that 20E functions in the IPCs. Our data indicate that EcR, Usp, E75 and DHR4 are required to promote IIS, whereas DHR3, and possibly also Ftz-f1, inhibit it. Based on these observations, we propose that 20E may regulate IIS similarly to how it regulates E biosynthesis in the PG. Here, the temporal cascade of nuclear receptors initially acts to promote E biosynthesis via early targets like E75 and DHR4, though later factors like DHR3 and, mainly, Ftz-f1, act to inhibit it (56). This serves to define the 20E peak as such and reduce 20E production using a self-regulated negative feedback. It would make sense, therefore, that 20E regulation of IIS in the IPCs would work similarly, as IPC-derived IIS is one of the strongest promoters of E biosynthesis during L3 development. Thus, 20E would serve initially as a feed-forward activator to stimulate IIS and, in turn, E biosynthesis. Then, once later nuclear receptors are expressed, they switch to negative feedback and inhibit IIS. Furthermore, regulation of the IPCs via these nuclear receptors could also involve other ligands, such as nitrous oxide (NO), juvenile hormone (JH) or others as yet unidentified, that interact with these nuclear receptors to exert their function in the IPCs, as they do in other tissues (57-59). Future studies will better define the temporal roles for these nuclear receptors in the IPCs and related neurons, and identify their respective ligands and interactors.

To conclude, we find that 20E bioactivation by *shade* is a nutrient-dependent process important in regulating both growth and developmental timing, which appears to depend on direct regulation in trachea and IPCs. Our work contributes important insights into how nutritional status is coordinated with maturation, via a complex integration of insulin and steroid signals.

## Materials & Methods

### Fly Stocks and Transgenes used in this Study

*Drosophila* stocks were reared at 25°C using standard yeast fly food recipe and rearing conditions. Stocks were ordered from the Bloomington *Drosophila* Stock Center (BDSC) or Vienna *Drosophila* RNAi Center (VDRC). The following stocks were used: Dilp2-GAL4^R^ (Provided by Dr. E. Rulifson; BDSC# 37516, described in (2)), *Dilp2-GAL4^215-1-1-1^* (46), *Dilp2-GAL4^96A08^* (47), *Dilp2-HA-FLAG* (provided by Dr. R Delanoue, described in (46,47)). To visualize IPCs, *Dilp2-GAL4* lines controlled expression of *UAS-sytGFP (35)*. Similarly, to perturb 20E signalling, *Dilp2-GAL4* lines controlled *UAS-RNAi* corresponding to genes of interest or *UAS-EcR^B1DN^* (36). *UAS-CYP18A1* (60) was a gift of Dr. K. Rewitz and *UAS-shade* (16) a gift of Dr. M. O’Connor. *Dahomey w*- flies, a gift of Dr. C Ribeiro, were used as a wildtype stock for positive controls in starvation experiments. RNAi lines used in this study include *EcR-RNAi* (BDSC# 50712), *Ftz-f1-RNAi* (BDSC# 33625, 27659), *E75-RNAi* (BDSC# 35780, 43231), *DHR3-RNAi* (VSC#: v106837), *DHR4-RNAi* (VSC#: v37066), *shade-RNAi* (VSC#: v17203, v106072) and *Usp-RNAi* (generated by Dr. C. Antonewski and provided by Dr. A. Andres and Dr. K. Lantz).

### Larval Starvation and *shade* Rescue Experiments

For starved/fed experiments, L3 larvae were staged and reared as above, then starved for 24h on 1% non-nutritive agar plates prior to collection and compared to fed L3 larvae from the same collection and identically handled. To control for crowding, embryos were divided with an equal density (50 embryos/agar plate). Fat bodies were carefully dissected in clean 1x PBS and snap-frozen before addition of QIAzol Lysis Reagent (QIAGEN Cat# 79306) and RNA extraction, cDNA synthesis and qPCR as described below.

For *shade* rescue experiments, *ppl*>*GFP* and *ppl*>*shade* L3 larvae were synchronously staged and reared at identical densities until L3 ecdysis before being subjected to 24h starvation or feeding on yeasted sucrose agar plates as described above. Whole L3 larvae were then collected and RNA extraction, cDNA synthesis and qPCR were performed as described below.

### Developmental Staging and Time to Pupariation

Staging of *Drosophila* larvae was done by collecting embryos during 2-3 hour intervals on yeasted, 20% sucrose 1% agar plates and visually scoring progression through instar stages at 25°C on a 12h light/dark cycle. Time to pupation was scored, where replicates of five to ten larvae were isolated just after L3 ecdysis and reared in normal conditions described above. The number of pupae were counted every 6 hours. The time to pupation of surviving larvae was plotted in GraphPad Prism software and compared to control siblings reared in identical nutritive conditions. The time to 100% pupation was compared between genotypes to statistically assess differences in developmental timing.

### Determination of Adult Fly Weight and Larval Growth Rates

After 3-7 days, experimental (*Dilp2-GAL4* > *UAS-EcR^B1DN^* or *UAS-RNAi, UAS-sytGFP*) and control siblings (*CyO; UAS-EcR^B1DN^* or *UAS-RNAi, UAS-sytGFP*) reared in identical environmental conditions were anesthetized using chloroform and weighed on a Mettler Toledo XS204 scale (Wet Weight; d = 0.1 mg). For the determination of larval growth rates, larvae were isolated after second instar (48h AEL) and weighed every 6 hours, starting in late L2, prior to third instar ecdysis (~72h AEL). Mean weight between experimental and control siblings was compared using a Student’s T-test with GraphPad software, and the change in adult weight between experimental and control genotypes were plotted. Larval weights were plotted as a function of time and a linear regression calculated to determine growth rates. The interaction terms of the regression lines for larval growth rates were then compared using GraphPad Prism software. Images of adult flies and larvae were taken using Olympus CellD software and figures were arranged in Adobe Photoshop CS4. For growth rate and weight experiments without internal control siblings (i.e. *ppl-GAL4*), animals were reared at controlled densities of ~50 larvae per plate.

For starvation weight experiments, larvae were weighed before and after 24h starvation, and the change in weight was plotted as a percent value. These experiments were also repeated weighing three larvae at a time to ensure accurate measurement and reproducibility. qPCR Analysis

### qPCR Analysis

Total RNA was isolated using TRI™ reagent (Invitrogen) and reverse transcription was performed on 1μg RNA using Transcriptor First Strand Synthesis kit (Roche). Total RNA from fat bodies was isolated using QIAzol lysis reagent optimized for fatty tissues (#79306). Primer sequences used are listed in Table S1. qPCR was performed on a Step-One-Plus using the SYBR Green detection system. Transcript levels were normalized using the geometric means of *RpS13 & Rp49* or *RpL3 & Rp49*. Mean ΔΦ values were statistically compared as described below. Relative quantitation of transcript levels to control genotypes were calculated using the ΔΔCt method and plotted with GraphPad Prism software to visualize expression differences.

### Immunohistochemistry and Nile Red staining

Adult and larval brains were dissected in 1x PBS prior to 30 minute fixation with 4% formaldehyde. For imaging fat bodies, synchronously timed larvae were inverted and fixed for 30 minutes in 4% formaldehyde prior to antibody or Nile Red staining.

Primary antibodies were incubated overnight at 4°C in PAXD. Antibodies used in this study were mouse α-GFP (University of Iowa, Developmental Studies Hybridoma Bank (DSHB) 8H11; 1:100), rabbit α-GFP (Life Technologies, Ref#: A6455; 1:1000), rabbit α-dFoxO, mouse α-EcR (common; 1:20) and mouse α-EcR-B1 (1:20; Ag10.2 & AD4.4, respectively, deposited to the DSHB by Carl Thummel and David Hogness), rabbit α-Dilp2 (1:1000; (2)), rabbit α-Dilp3, rabbit α-Dilp5, rabbit α-dFoxO (1:200, provided by Dr. P. Leopold), mouse α-Usp (1:20, provided by Dr. H. Krause), mouse α-E75B (1:20, provided by Dr. H. Krause), rabbit α-DHR3 (1:100, provided by Dr. H. Krause), rabbit α-Ftz-f1 (1:20, provided by Dr. Carl Wu).

Rabbit antibodies directed against the Dilp3 partial peptide sequence DEVLRYCAAKPRT and against the Dilp5 peptide sequence RRDFRGVVDSCCRKS were generated as a service by Thermo Fischer Scientific Inc. For immunostaining, α-Dilp3 antibodies were used at a dilution of 1:500 and α-Dilp5 antibodies used at a dilution of 1:200. Secondary antibodies used include goat α-mouse FITC & Cy3 and goat α-rabbit FITC & Cy3 (Jackson Immunoresearch). All secondary antibodies were used at a dilution of 1:200.

Fat body lipid droplets were stained with Nile Red (working concentration of 1:10,000) on inverted L3 larvae collected in 1x PBS and fixed for 20 minutes in PBT. Fat bodies were then removed and mounted in Vectashield^®^ Mounting Medium (Vector Laboratories Cat #H-1000).

Immunohistochemistry images were taken with an Olympus FluoView FV1000 confocal microscope and processed using ImageJ64 (1.6.0_65; FIJI) (61,62) and Photoshop CS4 software. In images of IPC morphology, contrast was enhanced to allow visualization of the thin proximal and medial neurites in addition to cell bodies and descending axons. Quantification of images was done using ImageJ64 software Measure tool for stainings and cell body sizes (largest cross-sectional area). To measure neurite branching complexity, the Plot Profile tool for thresholded (mean) stacks of IPCs was used from the midline outward to the longest, rightward proximal neurite.

### Analysis of larval feeding

For imaging food consumption, larvae were reared to L3 and transferred at 72h AEL to a yeasted 1 % sucrose agar plate, where both agar and yeast were supplemented with a red food colouring dye. Larvae were then imaged 3h after feeding and the presence of food dye in the digestive tract was used as a measure for feeding.

### Triglyceride Determination

Larvae were collected at ~96h AEL, snap-frozen and homogenized in 220 μL 0.05% Tween-20 in 1x PBS. Samples were heated to 70°C for 10 minutes to inactivate Lipase enzyme, and centrifuged. 25 μL of supernatant, 0.05% Tween-20 blank or a known concentration of Sigma Glycerol Standard (#G7793) was added to 200 μL of Infinity™ Triglyceride Reagent from Thermo Fischer Scientific (#981786) or Sigma Free Glycerol Reagent (#F-6428) and incubated for 60 minutes at 37°C. Absorbance was measured at 540nm on a TECAN M200 Pro spectrophotometer. Triglyceride measurements were normalized by total protein content, measured with the Pierce BCA protein assay kit at 562 nm, and visualized relative to control siblings reared in identical environmental conditions.

### Measurement of Circulating Dilp2-HA-FLAG

Measurement of hemolymph Dilp2 protein was performed as described in (45). Replicates of 7 larvae were gently bled on a depressed slide and 1 μL of hemolymph was collected. Samples were adhered to anti-FLAG (Sigma, F1804-50UG) coated 96-well plates together with anti-HA-Peroxidase (3F10; Roche, 12 013 819 001). Detection was performed using 1-step TMB Ultra ELISA substrate for 30 minutes at room temperature before absorbance values were measured at 450 nm. Mean circulating Dilp2-HA-FLAG was compared between samples and circulating Dilp2-HA-FLAG (Dilp2-HF) was plotted relative to *Dilp2-GAL4^215-1-1-1/+^* controls in an identical *w^1118^* genetic background (*w^1118^*) to *EcR^B1[DN]^* or *NR-RNAi* constructs. The standard used for determination was the synthetic peptide FLAG(GS)HA (DYKDDDDKGGGGSYPYDVPDYA) synthesized by LifeTein ^®^ LLC.

### Statistics

All statistics were conducted using the GraphPad Prism software. All statistics were performed on raw data after normalization, where applicable (i.e. qPCR). For assays comparing means, unpaired T-tests for pairwise comparisons (as in between *ppl* > *GFP* and *ppl* > *shade-RNAi* larvae) or one-way analysis of variance (ANOVA) tests for multiple comparisons were used. To test whether variance significantly differed between samples, an F-test was performed for pairwise comparisons and both Brown-Forsythe and Bartlett’s Tests performed for multiple comparisons. If the standard deviations between samples were significantly different, a T-test with Welch’s correction (Welch’s T-test) was performed for multiple comparisons, whereas the Geisser-Greenhouse correction was always applied on all one-way ANOVA analyses, as sphericity of the data was not assumed. When comparing means to a single control sample (as with *dilp2-GAL4^215-1-1-1/+^* and *dilp2-GAL4^215-1-1-1^* > *NR-RNAis*), Dunnett’s multiple comparisons test was used, and when comparing each mean to each other, Tukey’s multiple comparisons test was used.

For the comparison of growth rates, a linear regression was fit to raw weights distributed over defined time points. The slope of the curve defined the growth rate (gain in mass, x, over time, y). The R^2^ of the slope (Goodness-of-Fit) was calculated, and to test whether slopes are significantly different, an equivalent test to the analysis of covariance (ANCOVA) was performed to derive an F statistic and p-value (J Zar, Biostatistical Analysis, 2nd edition, Chapter 18, Prentice-Hall, 1984).

For all statistical analyses, we assumed a significance level of 0.05. For visualization of relative data, raw data was normalized to control samples either as a function of decimal (control = 1.0; i.e. qPCR analysis), percent (control = 100% i.e. TAG measurement) or change over time (Δ; control = 0.0%; i.e. weight and circulating Dilp2-HA-FLAG measurement).

## Abbreviations

Dilps: *Drosophila* insulin-like peptides
IPCs: Insulin-producing cells
IIS: Insulin/Insulin-like growth factor signalling
PG: Prothoracic Gland
20E: 20-hydroxyecdysone
L1-L3: First, Second, Third Larval Instar
FB: Fat Body
RNAi: RNA interference
qPCR: quantitative Polymerase Chain Reaction
h AEL: hours after egg laying
h AL3E: hours after third instar ecdysis

## Acknowledgments

The authors would like to thank Dr. Carl Thummel for providing the anti-DHR3 and anti-Usp antibodies, Dr. Henry Krause for the anti-E75B antibody, and Dr. Carl Wu for anti-ftz-f1 antibody. The authors also thank Dr. Andrew Andres and Kathryn Lantz for providing the *Usp-RNAi*, generated by Christopher Antonewski, Dr. Kim Rewitz for providing the *UAS-CYP18A1* line, Dr. Michael O’Connor for providing the *UAS-shade* line, Dr. Mark Metzstein for providing the *btl-GAL80* line and Dr. Pierre Leopold and Robert Harkness for critical feedback on this manuscript. K.B. is a predoctoral (“aspirant”) fellow of the Fonds Wetenschappelijk Onderzoek (FWO). This work was supported by VIB funding and FWO grants (G065408.N10 and G078914N) to P.C. The authors declare no conflict of interest.

## Author Contributions

K.B., J.C. and P.C. designed and interpreted the experiments, K.B. conducted the experiments, K.B. and M.W. conducted ELISA assays, K.B. and V.V. conducted *in situ* hybridization work and K.B., J.C., M.W. and P.C. wrote the paper.

